# Use of Transient Transfection for cGMP Manufacturing of eOD-GT8 60mer, a Self-Assembling Nanoparticle Germline-Targeting HIV-1 Vaccine Candidate

**DOI:** 10.1101/2022.09.30.510310

**Authors:** Vaneet K. Sharma, Vadim Tsvetnitsky, Sergey Menis, Evan T. Brower, Eddy Sayeed, Jim Ackland, Angela Lombardo, Thomas Hassell, William R. Schief

**Author notes:** **Correspondence**, Vadim Tsvetnitsky, William Schief.

## Abstract

We describe the current Good Manufacturing Practice (cGMP) production and subsequent characterization of eOD-GT8 60mer, a glycosylated self-assembling nanoparticle HIV-1 vaccine candidate germline-targeting priming immunogen. Production was carried out by transient expression in the human embryonic kidney 293 (HEK293) cell line followed by a combination of purification techniques. A large scale cGMP (200 L) production run yielded 354 mg of the purified eOD-GT8 60mer drug product material, which was formulated at 1 mg/mL in 10% sucrose in phosphate-buffered saline (PBS) at pH 7.2. The clinical trial material was comprehensively characterized for purity, antigenicity, glycan composition, amino acid sequence, aggregation, and by several safety-related tests during cGMP lot release. A comparison of purified products produced at 1 L scale and 200 L cGMP scale demonstrated consistency and robustness of the transient transfection upstream process and the downstream purification strategies. The cGMP clinical trial material is being tested in a phase 1 clinical trial (NCT03547245) and is currently stored at −80°C and on a stability testing program as per regulatory guidelines. The methods described here illustrate the utility of transient transfection for cGMP production of complex products such as glycosylated self-assembling nanoparticles.

## 1. INTRODUCTION

The goal of ending AIDS as a public health threat by 2030 envisaged for the world can only be realized by ensuring expanded and simplified HIV treatment and improved and effective use of interventions to prevent new infections (1). Despite steadily falling global human immunodeficiency virus (HIV) incidence, and reduction in the number of new infections to 1.7 million per year, there is widespread consensus that to eradicate HIV a vaccine is ultimately required (2).

Induction of broadly neutralizing antibodies (bnAbs), defined as antibodies capable of neutralizing diverse HIV isolates, is currently viewed as the central goal for HIV vaccine development. Results from the Antibody Mediated Prevention (AMP) trials provided evidence that passive immunization with a bnAb can protect against HIV infection (3, 4), consistent with similar studies in non-human primates (5). Germline-targeting vaccine design is one of several approaches currently under evaluation for generating bnAb responses to the highly variable HIV envelope (Env) protein (6–9). The germline-targeting strategy is based on the design of priming immunogens that induce responses from rare bnAb-precursor B cells. Sequential boosting with a series of immunogens then aims to guide antibody maturation to produce bnAbs (9–12).

VRC01-class bnAbs bind the CD4-binding site of HIV gp120 and include some of the most potent and highest breadth bnAbs known, making them important leads to guide vaccine design. At least three germline-targeting immunogens designed to prime VRC01-class responses are being manufactured for clinical testing: eOD-GT8 60mer (13–21), 426c core nanoparticles (22, 23), and BG505.SOSIP v4.1-GT1.1 (24, 25).

We describe the cGMP manufacture of eOD-GT8 60mer, a glycosylated self-assembling nanoparticle. In this molecule, the engineered outer domain of HIV gp120, germline-targeting version 8 (eOD-GT8) is fused to the C-terminus of a modified version of the enzyme lumazine synthase from the bacteria *Aquifex aeolicus* through a flexible glycine-serine linker. The fusion protein sequence contains 341 amino acids and has a molecular weight of 36618.1 Da. There are 8 paired cysteines and 10 N-linked glycosylation sites in the sequence. Upon secretion from mammalian HEK293 cells, the recombinant protein assembles into icosahedral nanoparticle structures (~30 nm diameter, ~3000 kDa molecular weight) composed of 60 identical subunits; the assembly process is driven by the inherent propensity for self-association of the lumazine synthase.

Although transient transfection is widely used in manufacturing of viral vectors and protein production to make non-clinical materials (26, 27), this approach is not common in the industry for generating protein products under cGMP conditions for clinical trials. For production of eOD-GT8 60mer, key components of the upstream processing included growth of suspension-adapted HEK293H cells, transient transfection components and conditions, feed strategy, and media and culture conditions. In the downstream processing, clarified harvest material was subjected to Benzonase treatment, viral inactivation (solvent/detergent treatment), purification by primary column chromatography on anion-exchange resin followed by polishing chromatography using ceramic hydroxyapatite resin, and nanofiltration-based virus reduction. Analytical characterization for the cGMP-produced eOD-GT8 60mer clinical trial material was based on established FDA requirements and ICH guidelines to determine safety, identity, concentration, purity, in-vitro potency and stability (28–30). Additional characterization included amino acid analysis (LC-MS/MS peptide mapping), N-linked glycan occupancy analysis (HILIC-FLD-MS), monosaccharide compositional analysis using RP-HPLC, particle size (DLS), and aggregation analysis using analytical ultracentrifugation (AUC). Finally, a nonclinical GLP repeat dose toxicity study was conducted in rabbits prior to a phase 1 clinical trial in healthy volunteers (NCT03547245).

## 2. MATERIALS AND METHODS

The eOD-GT8 60mer clinical trial material was manufactured at Paragon BioServices, Inc (Baltimore, MD), now part of Catalent Biologics under the name Paragon Gene Therapy. A cGMP facility was used to manufacture the clinical trial material using multiple unit operations, sterile techniques, and standard operation procedures. A working cell bank (WCB) of the suspension-adapted HEK293 cells, referred to as VRC293, was generated at SAFC (Carlsbad, CA) from a Master Cell Bank (MCB) generously provided by the National Institute of Health (NIH)/National Institute of Allergy and Infectious Diseases (NIAID)/Vaccine Research Center (VRC). The cGMP-qualified VRC293 cell line was generated from the HEK293H cell line at the VRC (31, 32). The VRC293 WCB was tested for absence of adventitious agents at release and for sterility and lack of mycoplasma contamination before cGMP process use. A vial of WCB was thawed and expanded for the manufacturing process. The plasmid DNA (pDNA) encoding eOD-GT8 60mer polypeptide chain was manufactured at Aldevron (Fargo, ND). Kifunensine, a chemically synthesized alkaloid, was manufactured under cGMP conditions by GlycoSyn (Graceville, New Zealand). Benzonase^®^ endonuclease enzyme (high-purity grade, 250 U/μL) was purchased from Millipore (Burlington, MA, USA). Polyethylenimide PEIpro -HQ^®^transfection reagent, a chemically synthesized polymer, was manufactured under cGMP conditions by PolyPlus-Transfection SA (Illkirch, France). Expi293 media was purchased from Thermo Fisher Scientific.

### 2.1 Transient Transfection and Upstream Process Development

The goal of the upstream process was to achieve high cell number, product titer and quality in the harvest (33, 34). VRC293 cells were expanded to grow in suspension in serum-free Expi293 media supplemented with 4 mM L-glutamine through a total of 8 passages via 1.0 L and 3.0 L shake flasks and 25 L Wave bioreactor (GE Healthcare) into a Biostat Cultibag STR 200 Plus (Sartorius Stedim Biotech) single-use production bioreactor. The cells in approximately 180 L working volume were transiently transfected with PEIpro-HQ^®^/ pDNA complex at cell density of *ca.* 10^6^ cells/mL (at 97.9% viability). To achieve high transfection efficiency, a solution containing 2:1 mass ratio of PEIPro-HQ^®^to pDNA was incubated for 12-15 minutes at ambient temperature and then added to VRC293 cells in OptiPRO FreeStyle serum-free medium (Gibco) at 37 °C. To fulfill regulatory requirements for using high-quality GMP source pDNA, quality control analytical testing was performed to ensure pDNA was sterile and of high purity and desired concentration. Additionally, tests were conducted to ensure that supercoiled pDNA was essentially free from genomic DNA, host proteins, residual RNA, and endotoxins (data not shown).

Kifunensine solution was added to the bioreactor after transfection to a final concentration of 14 μM to inhibit type-1 a-mannosidases in the endoplasmic reticulum (and likely also in the Golgi) and thereby maintain high-mannose glycoprotein levels during the transient transfection stage (17, 35, 36). Culture pH, temperature, dissolved oxygen, and agitation in the bioreactor were maintained post-transfection at pH 7.2 ±0.1, 37 ±2°C, 39 ±3% dissolved oxygen, and 75-90 rpm, respectively. During the run, cell density, cell viability, pH, and glucose and glutamine levels were monitored daily using a Vi-CELL™ XR Cell Viability Analyzer (Beckman Coulter, Inc., Fullerton, CA) and a BioFlex instrument (Nova Biomedical, Waltham, MA). Glucose and glutamine were supplemented after transfection as needed to avoid depletion. D-glucose (40%) was added on days 3 and 5, and L-glutamine (200 mM) was added on day 3, to maintain their respective concentrations (glucose ≥2 g/L and L-glutamine ≥2 mM). The cumulative amounts of glucose and glutamine added post transfection to the bioreactor were 4 kg and 3 kg, respectively. In the bioreactor, culture conditions such as impeller speed, pH, and temperature were also monitored daily. Seven days post transfection, the cells were separated from the conditioned supernatant by Unifuge (Pneumatic Scale Angelus, Stow, OH), a fully automated centrifuge system that uses a single-use insert inside a centrifuge bowl. Opticap XL capsule filters (EMD Millipore) were used for further cell separation and harvest clarification. The resulting clarified harvest was filtered through a 0.22 μm Millipore filter, and the product titer was determined to be 42.5 mg/L by a product specific enzyme-linked immunosorbent assay (ELISA).

### 2.2 Downstream Process

The downstream process, depicted in Figure 1, was focused on removing the process-derived and product-derived impurities, and reducing bioburden and endotoxin, while providing acceptable product yield (37, 38). The clarified harvest was concentrated approximately ten-fold by flat sheet tangential flow filtration (TFF) filters with a molecular cut-off 500 kDa (EMD Millipore). The flat sheet filters had a combined surface area of 4 m^2^. Concentrated harvest was treated with endonuclease to digest any remaining host cell and plasmid DNA to facilitate meeting a regulatory threshold for DNA contamination. Sucrose was added to 10% (w/v) final concentration before the storage of cell culture harvest at −80°C.

**Figure 1:**
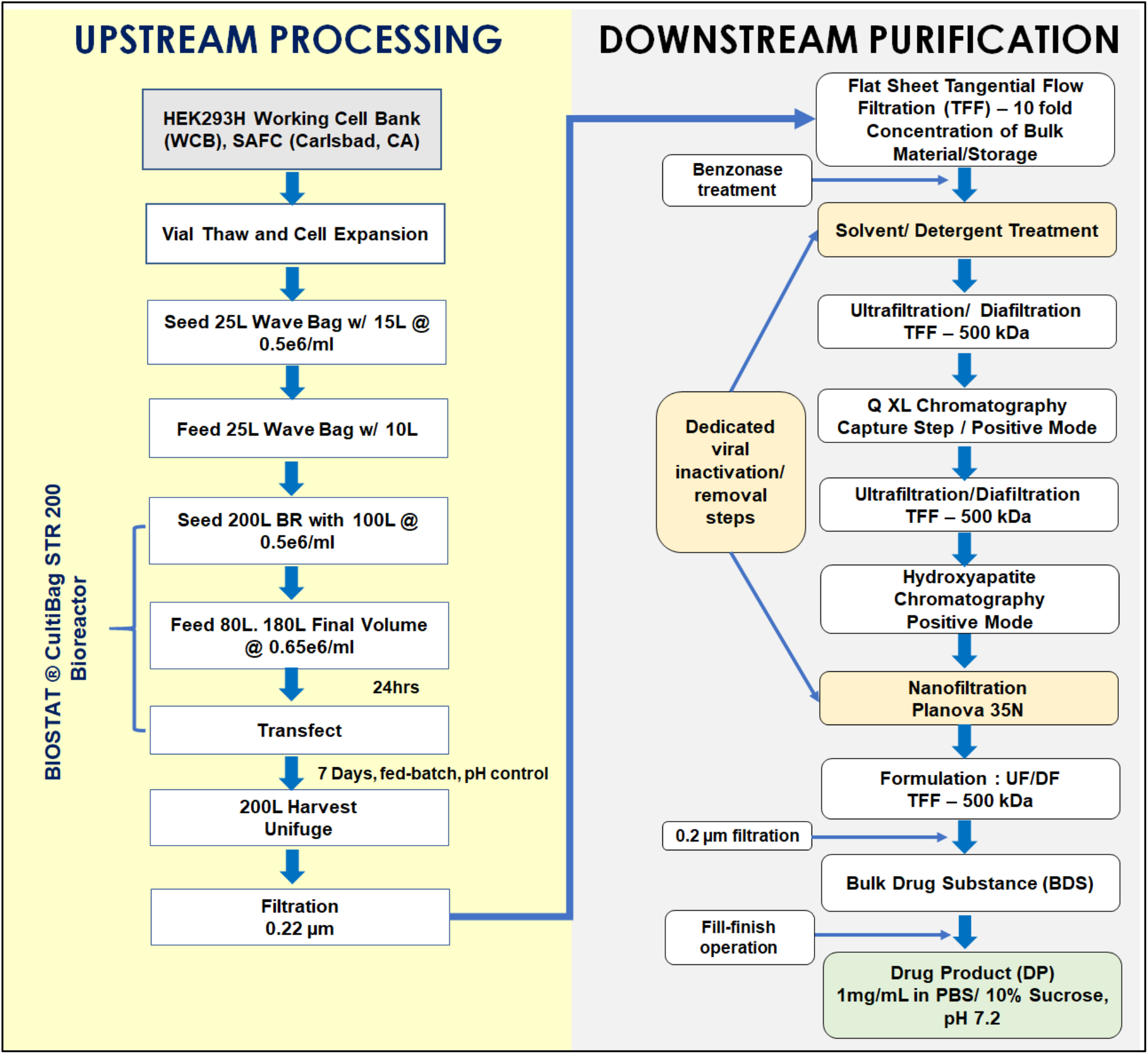
Overview of the various operations performed to transiently produce eOD-GT8 60mer from HEK293H cells and downstream processing to purify the nanoparticle.

Although HEK293 cells are not known to harbor known endogenous retroviruses, and adventitious agents were not detected in VRC92 MCB and WCB (results not shown), a solvent-detergent treatment step was incorporated in the manufacturing process out of an abundance of caution. The step was designed to inactivate enveloped viruses (39). Tri-n-butyl phosphate and Tween 80 were added to Benzonase-treated cell harvest to 0.3% and 1% final concentrations, respectively, and the solution was allowed to incubate for 16-20 hours at 2-8°C.

#### Q Sepharose XL Anion Exchange Chromatography

After endonuclease and solvent-detergent treatment, the clarified harvest was diafiltered into loading buffer (20 mM HEPES, 50 mM NaCl, pH 7.4) and loaded onto a 30-cm *(i.d.)* BPG™ 300 column (Amersham Biosciences Piscataway, NJ) packed with 1350 mL of Q-Sepharose XL resin (10 cm bed height) to perform anion exchange chromatography in bind-and-elute mode. Following the column washing, eOD-GT8 60mer captured by the resin was eluted isocratically using five column volumes of 20 mM HEPES and 250 mM NaCl buffer at pH 7.4 (flow rate 30 cm/h). In addition to providing enrichment of eOD-GT8 60mer, this chromatography step served to reduce the levels of contaminating DNA in the eluate. Even though the VRC293 cell line was derived from human tissue, residual host-cell protein and host-cell DNA in eOD-GT8 60mer product needed to be reduced to acceptable threshold values, typically below 100 μg/dose and 10 ng/dose (with DNA fragments not exceeding 200 base pairs), respectively, as per CBER guidelines, 2007.

#### Ceramic Hydroxyapatite (CHT) Column Chromatography

Ceramic hydroxyapatite (CHTTM, type 1, Bio-Rad, Hercules, CA), a crystalline mineral [(Ca5(PO4)3OH)2] with multimodal functionalities, was used in the bind-and-elute mode as a polishing step to remove aggregates and other product-related impurities. A CHT type 1 resin (80 μm particle size) packed in BPG™ 300 column (Amersham Biosciences Piscataway, NJ) was used to perform further eOD-GT8 60mer purification. Before the chromatography step, the packed column (6.3 L in volume) was equilibrated with 5 mM sodium phosphate, pH 7.2 equilibration buffer (≥ 4 CV). Prior to loading the Q Sepharose eluate onto the CHT column, the material was diafiltered into 5 mM sodium phosphate, pH 7.2 buffer. Under these conditions, the amino groups of eOD-GT8 60mer glycoprotein would be expected to bind to the phosphate ions of the mineral by a classical cation exchange mechanism, and carboxylic groups would be expected to bind to the Ca2+ ions of the resin by a combination of calcium metal affinity and anion exchange. Bound eOD-GT8 60mer glycoprotein was eluted from the CHT resin with a linear gradient from 0 to 100% of 500 mM sodium phosphate, pH 7 buffer over 10 column volumes at 9.4 L/hr. Fractions enriched with the target protein were pooled based on information obtained during the process development at small scale and an earlier engineering run performed at full 200 L production scale.

#### Viral Nanofiltration

The post-CHT pool was nanofiltered using single-use Planova 35N (35 nm nominal pore size, surface area 0.12 m^2^) membranes (Asahi Kasei Corporation, Tokyo, Japan) as a virus removing polishing step, which complemented the solvent/detergent viral inactivation operation performed earlier in the downstream purification. A formal viral clearance study demonstrated that for model viruses Planova 35N filtration provided ≥ 3.83 log clearance for xenotropic murine leukemia virus (XMuLV, 70-100 nm retrovirus) and 6.39 log clearance for the bovine viral diarrhea virus (BVDV, 40-70 nm pestivirus). After viral nanofiltration, the filtered CHT pool was TFF-formulated into a bulk Drug Substance (DS) by concentrating to 1.0 mg/mL of eOD-GT8 60mer and buffer-exchanging to 10 mM Na_2_HPO_4_, 2 mM NaH_2_PO_4_, 137 mM NaCl, 2.7 mM KCl, 10% sucrose at pH 7.4. A total of 354 mg eOD-GT8 60mer DS was purified from the starting 200 L cGMP batch (4.3 % process yield).

The formulated purified Drug Substance was filter-sterilized through a 0.22 μm filter (PVDF, Millipore, US) into a sterile single-use bag. The clinical trial material was filled at a 0.4 mL volume in 2 mL Type 1 glass vials with stoppers (13 mm stopper, Rubber with Flurotec, Afton Scientific) and sealed with sterile seals from Afton Scientific Corporation (Charlottesville, VA). Every sealed vial was subjected to visual inspection; vials were labeled and packaged into allocated fiberboard freezer boxes and kept at the intended storage temperature (−80°C).

Long-term stability studies for the cGMP batch of eOD-GT8 60mer clinical trial material were designed according to the ICH Q5A guideline. Stability of the clinical trial material stored at −80°C continues to be monitored over the 36-month period. Additionally, short-term stability studies were undertaken for liquid formulation stored at 5° ± 3°C (0, 24 hours) and at 25°C (0, 24 hours). At each time point testing is performed to evaluate appearance, pH, protein concentration, purity (SE-HPLC and SDS-PAGE), and relative potency. All the data generated to date (up to 36 months) demonstrated that the material continues to meet release specifications and no adverse trends in the stability-indicating parameters have been detected.

### 2.3. Analytical characterization

In the biopharmaceutical industry, comprehensive analytical characterization is required to support a product through development and cGMP manufacturing (28–30, 40). It is also a regulatory requirement to perform analytical testing on the cGMP manufactured clinical trial material (41, 42). Unlike most proteins such as monoclonal antibodies (~150 kDa) or other candidate HIV-1 vaccine immunogens (~120 - 400 kDa), eOD-GT8 60mer megadalton nanoparticle (~3000 kDa, ~30 nm diameter) posed significant characterization challenges to many traditional analytical quality control (QC) methods (43). QC procedures were performed on the manufactured eOD-GT8 60mer nanoparticle to confirm its identity, determine protein concentration and purity, establish in-vitro potency and measure the nanoparticle size. Additionally, to help assure safety for clinical trial participants, the clinical trial lot was tested to quantify host cell residual impurities (host cell proteins and host cell DNA), measure bioburden and bacterial endotoxin, determine subvisible particulate matter, and confirm sterility.

Although a specification-based analytical control strategy was established to ensure the quality and safety, the complex nature and composition of eOD-GT8 60mer nanoparticle could lead to product variation influenced by upstream conditions or downstream purification processes. To address this possibility, high-resolution analytics were carried out to physiochemically characterize the eOD-GT8 60mer nanoparticle (28, 40, 44–47). Overlapping information derived from the orthogonal analytical assays not only facilitated a better understanding of the structure and composition of eOD-GT8 60mer clinical trial material, but also assured product safety.

As the transient transfection-based process was developed at 2.0 L scale, optimized at 10.0 L volume, and further scaled-up to the 200 L cGMP clinical batch, the product critical quality attributes (CQAs) were established and monitored during scale-up to demonstrate product consistency. A combination of biophysical, immunochemical, and physicochemical methods was used to perform comparability testing to assess antigenicity, three-dimensional structures, post-translational modifications, and protein aggregation. A systematic characterization performed throughout the project validated reliable and reproducible scale-up, and batches for the eOD-GT8 60mer nanoparticle produced at different scale demonstrated near identical CQAs.

A sandwich ELISA was developed to characterize the potency of eOD-GT8 60mer glycoprotein (Figure 2a). This assay measured the binding of the germline (GL) VRC01 antibodies to the eOD-GT8 60mer nanoparticle. ELISA was performed using the 96-well Maxisorp ELISA plates (Thermo Fisher Scientific), coated over-night with 100 μL/well of 4 μg/mL GL-VRCO1 Fab (produced in-house at TSRI, San Diego, CA). The plates were incubated at 2-8°C for 18 ± 2 hours, followed by washing each plate three times with the wash buffer (PBS, 0.05% Tween 20) and subsequently blocking at 25°C for 45 ± 2 minutes while rotating at 180 RPM with 200 μL of BSA Blocking Buffer (PBS, 0.05% Tween 20, 2% BSA). After incubating with serially diluted samples containing eOD-GT8 60mer, 100 μL of 100 ng/mL of GL-VRC01 antibody solution was added to each well and incubated at 25°C for 45 ± 2 minutes for at 180 RPM. This was followed by adding 100 μL of diluted secondary HRP conjugated goat anti-human IgG Fcγ antibody to each well and incubating at 25°C for 45 ± 2 minutes for at 180 RPM. After three washes with wash buffer, 100 μL TMB substrate (Sigma Aldrich) was added to develop the chromogenic signal (10 minutes in the dark, 25°C). The reaction was stopped with 100 μL of 2N Sulphuric acid, and the absorbance at 450 nm was measured using the using a Spectramax microplate reader. The results were analyzed using Softmax Pro software (Molecular Devices LLC, Sunnyvale, CA, USA). In vitro relative potency was expressed as a percentage and defined as a ratio of protein concentration determined by ELISA to protein concentration determined by measuring absorbance at 280 nm. For all tested eOD-GT8 60mer samples, the potency was determined to be between 89-100 % (Table 1).

**Figure 2:**
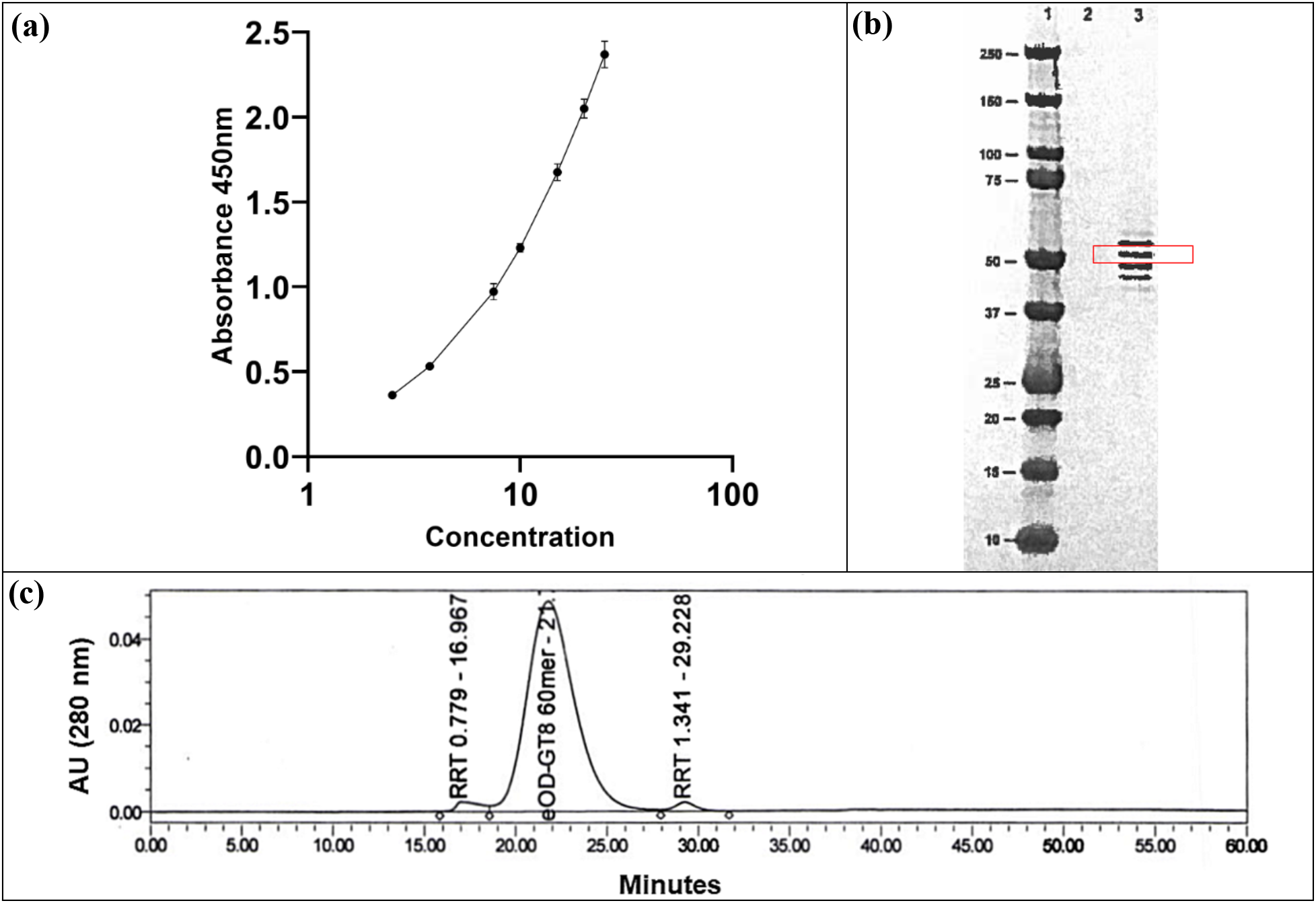
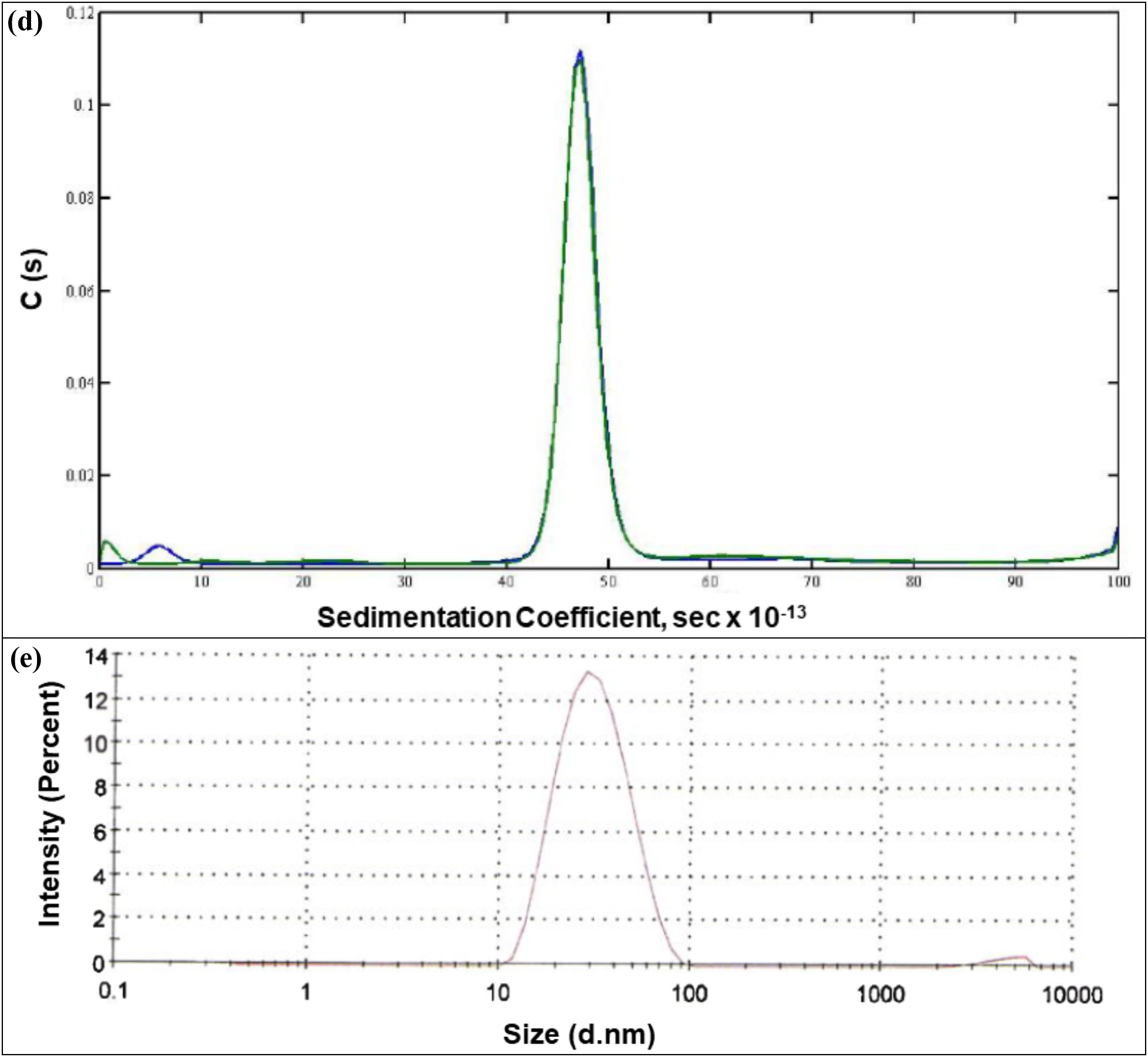
Analytical Characterization of the eOD-GT8 60mer. (a) Sandwich ELISA standard curve for the in-vitro potency analysis. (b) N-terminal sequencing by Edman degradation. The band enclosed in red rectangle was excised. (c) SE-HPLC analysis to monitor product purity. (d) AUC-SV analysis to determine peak distribution. (e) Dynamic Light Scattering (DLS) to determine average particle size via size distribution by intensity.

**Table 1:**
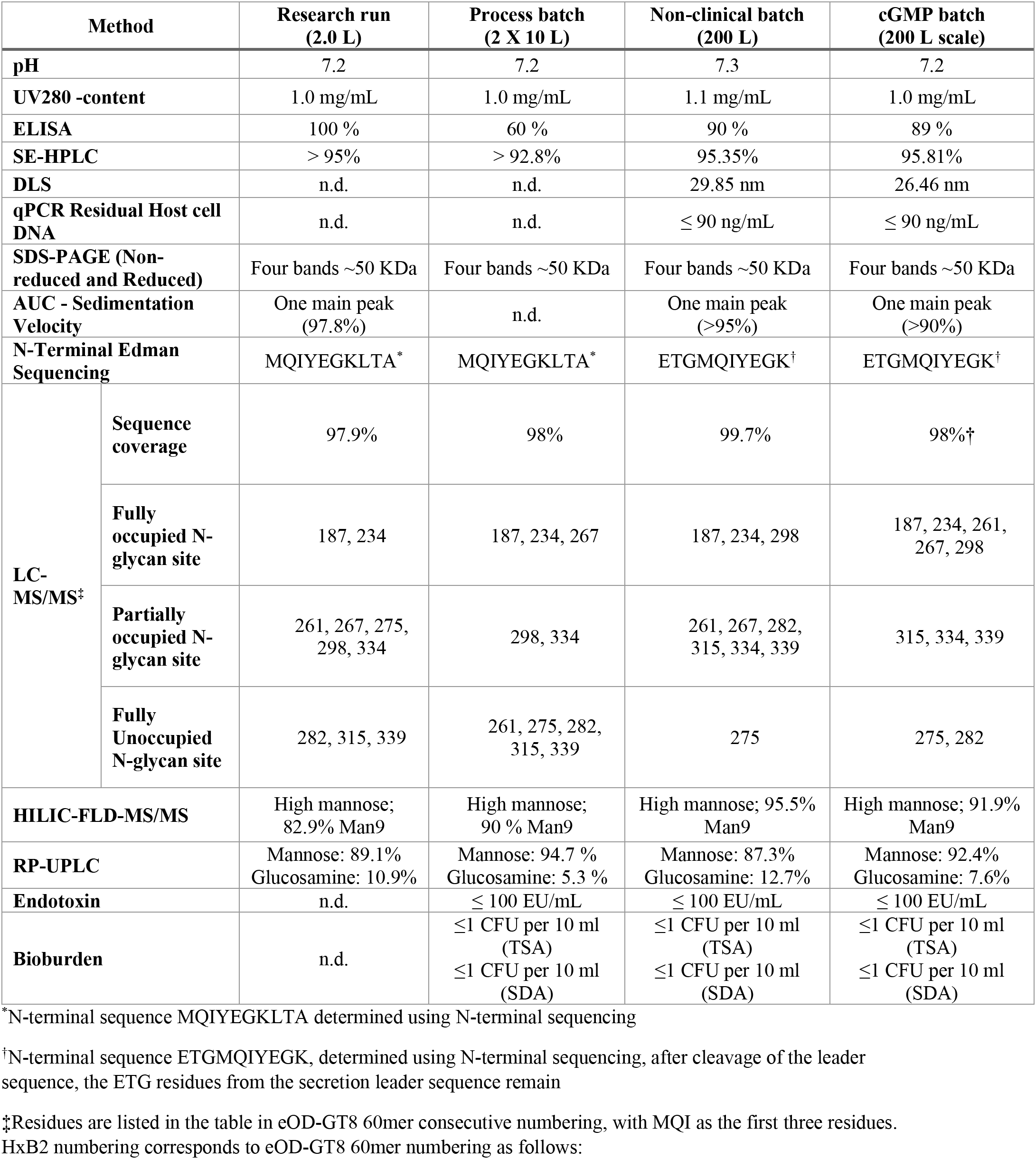
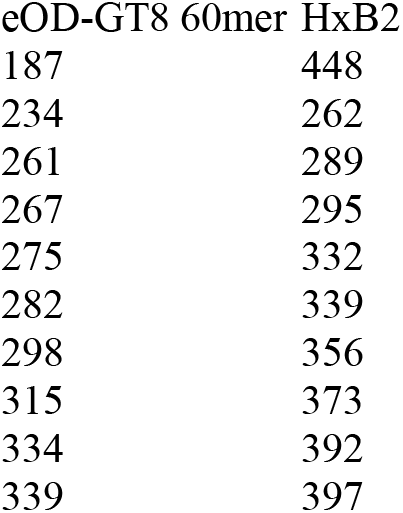
Comparison of eOD-GT8 60mer nanoparticle product characteristics from research (2.0 L), process development (10L) and large scale (200 L) production runs using transient transfection in HEK293H cells.

| Method |  | Research run<br>(2.0 L) | Process batch<br>(2 X 10 L) | Non-clinical batch<br>(200 L) | cGMP batch<br>(200 L scale) |
| --- | --- | --- | --- | --- | --- |
| **pH** |  | 7.2 | 7.2 | 7.3 | 7.2 |
| **UV280 -content** |  | 1.0 mg/mL | 1.0 mg/mL | 1.1 mg/mL | 1.0 mg/mL |
| **ELISA** |  | 100 % | 60 % | 90 % | 89 % |
| **SE-HPLC** |  | > 95% | > 92.8% | 95.35% | 95.81% |
| **DLS** |  | n.d. | n.d. | 29.85 nm | 26.46 nm |
| **qPCR Residual Host cell DNA** |  | n.d. | n.d. | ≤ 90 ng/mL | ≤ 90 ng/mL |
| **SDS-PAGE (Non-reduced and Reduced)** |  | Four bands ~50 KDa | Four bands ~50 KDa | Four bands ~50 KDa | Four bands ~50 KDa |
| **AUC - Sedimentation Velocity** |  | One main peak (97.8%) | n.d. | One main peak (>95%) | One main peak (>90%) |
| **N-Terminal Edman Sequencing** |  | MQIYEGKLTA* | MQIYEGKLTA* | ETGMQIYEGK† | ETGMQIYEGK† |
| **LC-MS/MS‡** | **Sequence coverage** | 97.9% | 98% | 99.7% | 98%† |
|  | **Fully occupied N-glycan site** | 187, 234 | 187, 234, 267 | 187, 234, 298 | 187, 234, 261, 267, 298 |
|  | **Partially occupied N-glycan site** | 261, 267, 275, 298, 334 | 298, 334 | 261, 267, 282, 315, 334, 339 | 315, 334, 339 |
|  | **Fully Unoccupied N-glycan site** | 282, 315, 339 | 261, 275, 282, 315, 339 | 275 | 275, 282 |
| **HILIC-FLD-MS/MS** |  | High mannose; 82.9% Man9 | High mannose; 90 % Man9 | High mannose; 95.5% Man9 | High mannose; 91.9% Man9 |
| **RP-UPLC** |  | Mannose: 89.1%<br>Glucosamine: 10.9% | Mannose: 94.7 %<br>Glucosamine: 5.3 % | Mannose: 87.3%<br>Glucosamine: 12.7% | Mannose: 92.4%<br>Glucosamine: 7.6% |
| **Endotoxin** |  | n.d. | ≤ 100 EU/mL | ≤ 100 EU/mL | ≤ 100 EU/mL |
| **Bioburden** |  | n.d. | ≤1 CFU per 10 ml (TSA)<br>≤1 CFU per 10 ml (SDA) | ≤1 CFU per 10 ml (TSA)<br>≤1 CFU per 10 ml (SDA) | ≤1 CFU per 10 ml (TSA)<br>≤1 CFU per 10 ml (SDA) |

\*N-terminal sequence MQIYEGKLTA determined using N-terminal sequencing
†N-terminal sequence ETGMQIYEGK, determined using N-terminal sequencing, after cleavage of the leader sequence, the ETG residues from the secretion leader sequence remain
‡Residues are listed in the table in eOD-GT8 60mer consecutive numbering, with MQI as the first three residues. HxB2 numbering corresponds to eOD-GT8 60mer numbering as follows:

|  |  |
| --- | --- |
| 187 | 448 |
| 234 | 262 |
| 261 | 289 |
| 267 | 295 |
| 275 | 332 |
| 282 | 339 |
| 298 | 356 |
| 315 | 373 |
| 334 | 392 |
| 339 | 397 |

The eOD-GT8 60mer amino acid sequence was confirmed using Agilent 1290 Infinity II HPLC coupled to Agilent 6550 QTOF Mass Spectrometer. A Waters XBridge CSH C18 column (130 Å, 1.7 μm, 2.1 × 150 mm) was used to perform chromatographic separation. The mobile phases used were 0.1% formic acid (mobile phase A) and 0.1% formic acid in 100% acetonitrile (mobile phase B). Bottom-up proteomics techniques (enzymatic digestion followed by MS analysis of proteolytic peptide mixtures) were used to perform peptide mapping. The eOD-GT8 60mer glycoprotein was reduced and carboxymethylated followed by lysine-C (Lys-C)/ trypsin sequential digestion followed by treatment with Glutamyl endopeptidase (Glu-C). The MS data were acquired in positive ionization mode over the range between 225 and 2500 m/z and MS/MS fragmentation data was acquired over the range between 100 m/z and 3000 m/z. The total ion chromatograms were analyzed using Agilent MassHunter software, and sequence matching was carried out using Agilent BioConfirm software. For all tested eOD-GT8 60mer samples, over 95% amino acid sequence was positively mapped including the N- and C-termini sequence and similar PTMs (deamidation, oxidation, pyroglutamylation, and N-linked glycosylation) were identified (Table 1).

N-terminal amino acid sequencing for the eOD-GT8 60mer was performed using the Procise Protein Sequencing System (Applied Biosystems, CA, USA). The protein band corresponding to the eOD-GT8 60mer (~50 kDa) was excised from SDS-PAGE gel (Figure 2b), transferred to a clean Eppendorf tube and subjected to 10 cycles of Edman degradation as described by Tempst and colleagues (48). The N-terminal amino acid sequence was determined to be ETGMQIYEGK for the non-clinical (200L) and cGMP (200L) batch (Table 1). The ETG residue on the N-terminal side for the non-clinical and cGMP batch remained from the secretion leader sequence after cleavage of the leader sequence. Additional quality attributes critical to eOD-GT8 60mer structure were evaluated and are listed in Table 1. Size exclusion-high pressure liquid chromatography (SEC-HPLC) was performed to monitor product purity. SEC-HPLC was carried out using an Alliance e2695 HPLC system (Waters Co., Milford, MA, USA) equipped with a Superose 6 Increase 10/300 GL column (GE Healthcare Life Sciences) and a 2998 photodiode array (PDA) detector. 100 μg of sample was directly injected onto the column at 25°C and a mobile phase consisting of 10mM HEPES 150mM NaCl pH 7.4, was used. The flow rate was 0.5 mL/min, and main peak of 60mer and impurities were detected under native conditions with UV detection at 280 nm wavelength. Data were acquired and processed by the Empower™3 (Waters) software.

The size distribution analysis of eOD-GT8 60mer samples was carried out by sedimentation velocity analytical ultracentrifugation (AUC-SV) on a Beckman Optima XL-A Ultracentrifuge equipped with a four-hole rotor (47). A sample held within one of the two channels in a 12-mm double-sector Epon centerpiece was sedimented at a moderate speed (20,000 RPM). All the tested samples on SEC-HPLC and AUC showed no aggregates and only one main peak corresponding to the eOD-GT8 60mer (Figure 2c, 2d) without any evidence of particle disassembly into smaller pentameric or monomeric components. The average particle size of eOD-GT8 60mer was measured as 29 nm (Figure 2e) by dynamic light scattering (DLS) with the Zetasizer NanoZS (Malvern, Southborough, MA)(46).

N-linked glycan profiling was performed by Charles River Laboratories using hydrophobic interaction liquid chromatography coupled to mass spectrometry (HILIC-FLD-MS/MS) to determine the type of N-linked glycans present on the eOD-GT8 60mer (Figure 3a). A two-step N-glycan release procedure was implemented to ensure the completeness of the deglycosylation reaction. Samples were first treated by non-denaturing release overnight with N-glycanase (Prozyme) at 37°C and, in a second step, a denaturation buffer containing SDS was applied prior to another overnight N-glycanase digestion. The released glycans were 2-AB labeled (Sigma-Aldrich) and quantified using HILIC-FLD equipped with fluorescence detector (excitation and emission wavelengths of 330 nm and 420 nm, respectively). The eluting N-linked glycans were further characterized using a high-resolution Agilent ESI-QTOF mass spectrometer model 6550. Man9 *[(Man)9(GlcNAc)2]* was the most abundant N-linked glycan for all the tested sample (Figure 4, Table 1). A high resolution LC-MS/MS peptide mapping based method was used to determine the sequence coverage and N-linked glycosylation site identification. The sample was treated with PNGase-F to remove N-glycans and digested with the trypsin coupled with Lys-C followed by GluC. The occupancy of N-glycans at each potential site was determined from the MS1 and MS2 data extracted from MS raw files processed using the Agilent Masshunter software and BioConfirm tools. The MS2 analysis reduced misinterpretation of glycosylation site occupancy by MS1 data alone from potential deamidation of Asn and/or Gln residue. LC-MS/MS analysis assessed de-N-glycosylation producing Asn to Asp conversion and a corresponding mass shift of +0.98 Da to identify the potential N-glycan sites as occupied, partially occupied, or unoccupied, using a method similar to but less quantitative in measuring occupancy than the methods described by Paulson and colleagues (44, 45, 49). Apart from some minor differences all the tested samples (Table 1) exhibited similar sequence coverage and N-glycan occupancy. Five N-glycan sites were fully occupied, three were partially occupied, and two were not occupied.

**Figure 3:**
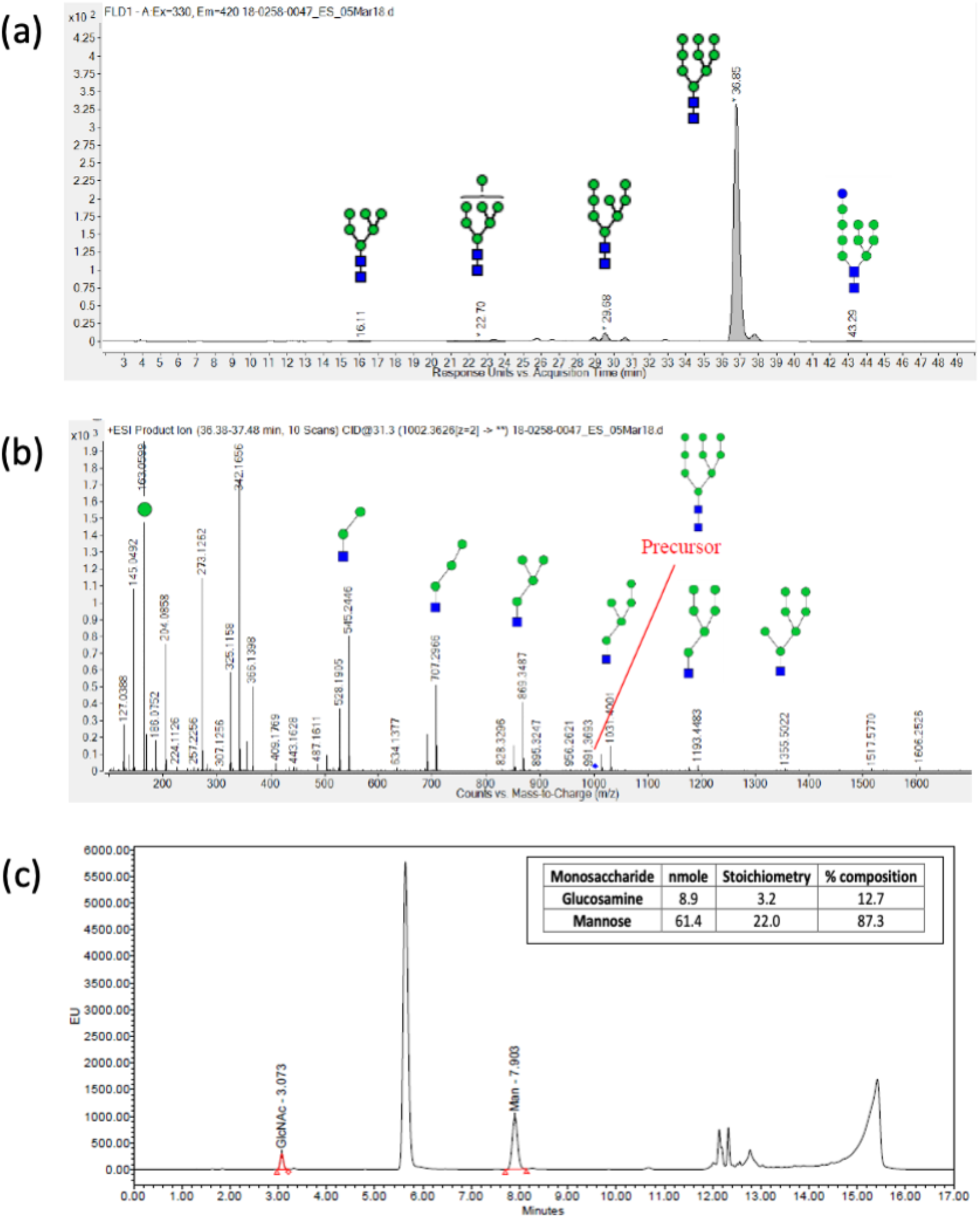
N-linked glycans analysis of eOD-GT8-60mer sample. (a) Released N-glycans analysis using UPLC-HILIC-FLD. Man9 (95.5%) was determined to be the most dominant N-linked glycans. (b) MS/MS of Man9 using Q-TOF/MS. (c) Quantitative monosaccharide composition analysis using RP-HPLC.

**Figure 4:**
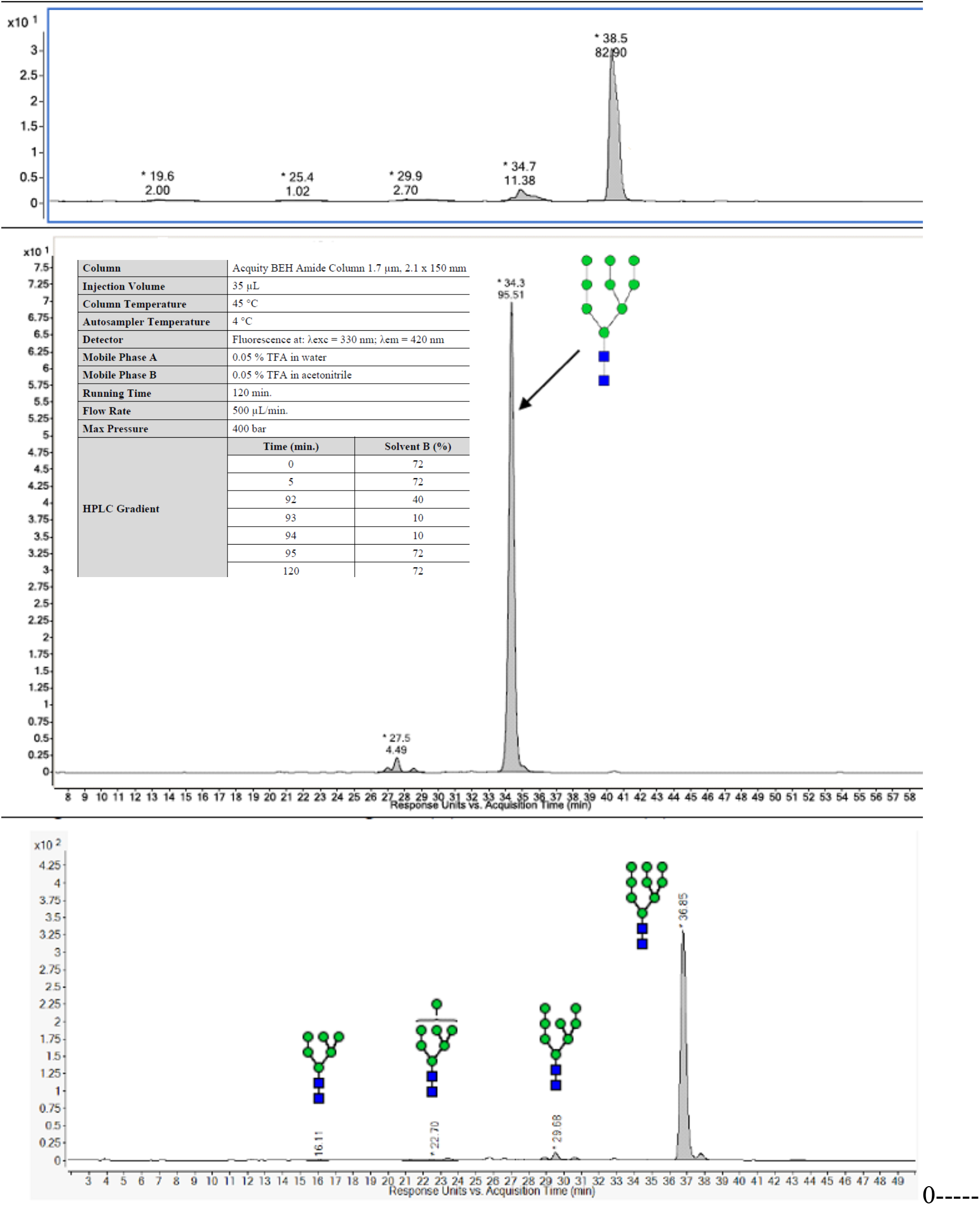
N-linked glycan comparison for the process (10L), non-clinical batch (200 L) and clinical batch (200 L) scale runs. The difference in the retention time is due to different HPLC used. The run conditions were identical, detailed in the in the middle panel. The N-linked glycans structure was confirmed by MS/MS.

In addition to N-linked glycan analysis, monosaccharides analysis was also performed to confirm the composition of the sugars in the eOD-GT8 60mer glycoprotein (50). The test sample was hydrolyzed using trifluoroacetic acid (TFA) and then derivatized using fluorescent 2-aminobenzoic acid (2-AA) prior to reversed phase chromatographic analysis (RP-HPLC). The separated monosaccharide derivatives were detected by their fluorescence intensity on an ultra-high-performance liquid chromatography (UPLC-FLD) linked with a fluorescence detector (excitation and emission wavelengths of 360 nm and 425 nm, respectively). In all tested samples, mannose and glucosamine were the only two sugars detected (Figure 3b, Table 1).

The multifaceted eOD-GT8 60mer nanoparticle characterization revealed similar product characteristics among different batches. The analytical testing results for the batches produced at a different scale (i.e., 1.0 L research run, 20 L process run, 200 L engineering run, and 200 L cGMP batch) for the specified attributes, are summarized in Table 1.

### 2.4. Nonclinical safety assessment

The objectives of this study were to determine the potential toxicity and local tolerance of eOD-GT8 60mer/AS01_B_ vaccine when given by intramuscular (IM) injection, and eOD-GT8 60mer when given by intramuscular or subcutaneous (SC) injection. The vaccine products were given once every two weeks for 4 doses to New Zealand White rabbits. The following parameters and end points were evaluated in this study: clinical signs, dermal scores, body weights, body weight gains, food consumption (males only), ophthalmology, body temperature, clinical pathology parameters (hematology, coagulation, clinical chemistry, and c-reactive protein) and immunogenicity (antibody analysis). Two (Day 45) and 21 (Day 64) days after the last injection gross necropsy findings and organ weights were collected with subsequent histopathologic examinations.

The intramuscular injection of OD-GT8 60mer/AS01B vaccine, was well tolerated and was only associated with minor disturbances in clinical pathology, higher c reactive protein levels, and findings of mixed inflammatory cell infiltrate at the administration sites. The intramuscular or subcutaneous injection of eOD-GT8 60mer, was well tolerated and was only associated with findings of mixed inflammatory cell infiltrate at the administration sites. After a 3-week treatment free period, the findings were no longer recorded or were at lower incidence and severity. There were no findings that were considered to be adverse.

## 3. DISCUSSION

Transient transfection is a common laboratory technique to quickly produce small amounts of protein from mammalian cell culture. While it is widely used now at larger scale to produce clinical supplies of viral vectors for gene therapy, production of clinically relevant proteins using transient transfection have not been reported. Here we describe a reproducible and scalable transient transfection-based process for GMP manufacturing of an HIV-1 vaccine priming candidate with complex molecular features (glycosylated, self-assembling nanoparticle), and we detail an analytical testing strategy for product characterization and release.

It is important, but challenging, to maintain comparable process performance for each unit operation upon scale up. During transient transfection, plasmid DNA must be mixed with a transfection reagent and then added to the bioreactor. Fluid dynamics in different bioreactor volumes are a concern, as there is a narrow window of time in which mixing and subsequent addition to the cells should be achieved to maintain consistent transfection efficiency and production yield. Here we demonstrate that transient transfection of HEK293 cells with PEI as transfection reagent at 2, 10 and 200 L scale provided very similar product titers in the harvest and indistinguishable quality of the final product after downstream purification. Process conditions employed for the engineering run conducted at full 200L scale were then faithfully reproduced for the cGMP run at the same scale, resulting in product batches with the same product characteristics.

In the upstream process, the consistency of production was achieved by identifying conditions that minimized cell clumping and ensuring optimal feeding to maintain high HEK293 cell viability. The downstream process allowed for efficient purification of the megadalton nanoparticle from process- and product-related impurities. eOD-GT8 60mer bound efficiently to anion exchange resin in the first chromatography step, allowing for significant enrichment in the eluate, and the subsequent hydroxyapatite column provided efficient separation from remaining impurities. The two-column process was supplemented with viral inactivation and removal steps. Additionally, we took advantage of the large molecular weight of the product and utilized large pore size TFF membrane for buffer exchange and concentration steps, while achieving some additional polishing.

We utilized an extensive analytical toolbox and sophisticated techniques in order to better understand the product characteristics and to obtain baseline data for future manufacturing batches. In addition to confirming identity, potency and safety of the clinical product, we demonstrated consistent average size of eOD-GT8 60mer nanoparticles and lack of aggregation. For consistency with pre-clinical experiments, Kifunensine was used to ensure that the glycans would be high mannose. N-linked glycan profiling of the product confirmed the predominance on Man9 among eluted sugars. N-linked glycan occupancy analyses showed that five of the ten N-linked glycosylation sites were partially or fully unoccupied. Pre-clinical material, produced by transient transfection in 293 cells and purified using lectin affinity followed by size exclusion chromatogrphy, had seven N-glycan sites partially or fully unoccupied (19) but nevertheless performed well in multiple mouse models (14, 15, 17, 20, 21, 51, 52). Therefore, we viewed underoccupancy of glycans as unlikely to hamper performance in human clinical testing. Whether glycan occupancy would be improved by production from a stable cell line is not known. As quality control analytical testing alone does not ensure product safety, a non-clinical safety assessment of the eOD-GT8 60mer produced at 200 L batch scale was performed for increase assurance of safety. There were no findings considered to be adverse.

The pursuit of a safe and effective HIV vaccine is a significant public health priority. The work described here supported the clinical testing of a new immunogen, eOD-GT8 60mer, and a new vaccine concept, germline targeting. A randomized, double-blind, placebo-controlled dosage-escalation Phase 1 study is being conducted to assess the safety, tolerability, and immunogenicity of eOD-GT8 60mer vaccine, adjuvanted with AS01_B_, in up to 48 healthy adult HIV-negative volunteers (ClinicalTrials.gov Identifier: NCT03547245). Experimental medicine clinical trial studies to evaluate new HIV vaccine candidates are novel, and manufacturing these products in accordance with cGMP is challenging. The recovery of highly pure and stable eOD-GT8 60mer from a cGMP process based on transient transfection of HEK293 cells provides a potential template for cGMP manufacture of other immunogens. Relatively small amounts of produced Drug Product were sufficient for lot release and stability testing, and, given low clinical dose, provided enough material for potentially several Phase 1 studies. The reliability and robustness built through comprehensive analytical characterization have established procedures that are adaptable to successful cGMP campaigns for future immunogen-based HIV vaccine projects.

## ACKNOWLEDGMENTS

This work was supported by the Bill and Melinda Gates Foundation Collaboration for AIDS Vaccine Discovery grants to the IAVI Neutralizing Antibody Consortium (INV-007522 and INV-008813). The cGMP manufacturing work was funded by a grant to IAVI from the Bill & Melinda Gates Foundation. We are grateful to Pervin Anklesaria (BMGF) for input and support. IAVI is grateful to the NIH/DAIDS/VRC for their generous donation of the VRC293 working cell bank. We thank the project team from Paragon BioServices, Inc (Baltimore, MD), now part of Catalent Biologics under the name Paragon Gene Therapy, for the process development and cGMP manufacturing. We thank Diane M. Kubitz and her team for producing and testing the required antibodies for this program at The Scripps Research Institute Antibody Core for antibody production. We acknowledge Sam Pallerla, Roslyn Platt, Kristen Syvertsen, Natasha Williams from IAVI, and Nicole Yates, LaTonya Williams, Kelli Greene and Hongmei Gao from CAVIMC-Montefiori, Tomaras group at Duke Human Vaccine Institute for their expert technical assistance. We thank Liwei Cao, Jolene K. Diedrich, John R. Yates III, and James C. Paulson for mass spectrometry-based site-specific glycan profiling of samples during method development prior to the work described here. We thank GlaxoSmithKline for their support in providing AS01B adjuvant.

